# Isoform-specific N-linked glycosylation of voltage-gated sodium channel α-subunits alters β-subunit binding sites

**DOI:** 10.1101/2024.05.20.594950

**Authors:** Christopher A. Beaudoin, Manas Kohli, Samantha C. Salvage, Hengrui Liu, Samuel J. Arundel, Samir W. Hamaia, Ming Lei, Christopher L.-H. Huang, Antony P. Jackson

**Affiliations:** Department of Biochemistry, Hopkins Building, University of Cambridge, Tennis Court Road, Cambridge, CB2 1QW, United Kingdom; Institute of Cancer Research, London, United Kingdom SW7 3RP, United Kingdom; Department of Pharmacology, University of Oxford, Oxford, OX1 3QT, United Kingdom.; Department of Physiology, Development and Neuroscience, University of Cambridge, Cambridge, CB2 3EG, United Kingdom

**Keywords:** Voltage-gated sodium channel, N-linked glycosylation, Nav1.5, Nav1.8, Nav β-subunit, Nav β1, ephaptic conduction

## Abstract

**Highlights:**

- Three N-linked glycosylation sites conserved among all Nav channels
- Glycan modelling and molecular dynamics simulations highlight 3D landscape
- Unique Nav1.5 N-linked glycans may prevent binding to Ig-domains of β1 and β3
- Unique Nav1.8 N-linked glycan may prevent binding to Ig-domains of β2 and β4
- N-linked glycans likely contribute to supra-molecular clustering of Nav channels

Voltage-gated sodium channel α-subunits (Nav1.1-1.9) initiate and propagate action potentials in neurons and myocytes. The Nav β-subunits (β1-4) have been shown to modulate α-subunit properties. Homo-oligomerization of β-subunits on neighboring or opposing plasma membranes has been suggested to facilitate cis or trans interactions, respectively. The interactions between several Nav channel isoforms and β-subunits have been determined using cryogenic electron microscopy (cryo-EM). Interestingly, the Nav cryo-EM structures reveal the presence of N-linked glycosylation sites. However, only the first glycan moieties are typically resolved at each site due to the flexibility of mature glycan trees. Thus, existing cryo-EM structures may risk de-emphasizing the structural implications of glycans on the Nav channels. Herein, molecular modelling and all-atom molecular dynamics simulations were applied to investigate the conformational landscape of N-linked glycans on Nav channel surfaces. The simulations revealed that negatively-charged sialic acid residues of two glycan sites may interact with voltage-sensing domains. Notably, two Nav1.5 isoform-specific glycans extensively cover the α-subunit region that, in other Nav channel α-subunit isoforms, corresponds to the binding site for the β1-(and likely β3-) subunit immunoglobulin (Ig) domain. Nav1.8 contains a unique N-linked glycosylation site that likely prevents its interaction with the β2 and β4-subunit Ig domain. These isoform-specific glycans may have evolved to facilitate specific functional interactions, for example by redirecting β-subunit Ig-domains outwards to permit cis or trans supra-clustering within specialized cellular compartments such as the cardiomyocyte perinexal space. Further experimental work is necessary to validate these predictions.

## 1. Introduction

Voltage-gated sodium (Nav) channels initiate and propagate the rising phase of the action potential in electrically-excitable cells, such as neurons and myocytes (Yu and Catterall, 2003). Nav channels are transmembrane proteins that consists of an ion-selective α-subunit (molecular mass ∼ 250 kDa) and associated regulatory β-subunits (molecular mass ∼ 30 kDa). There are nine structurally distinct Nav channel α-subunit isoforms (Nav1.1-1.9), which are relatively tissue-specific: e.g. Nav1.1-1.3 and Nav1.6 are largely found in the central nervous system, Nav1.4 is found in skeletal muscle, Nav1.5 is primarily expressed in cardiac muscle, and Nav1.7-1.9 are considered to be specific to the peripheral nervous system (Clare et al., 2000). However, several exceptions in tissue-specific Nav channel expression have been discovered. For example, Nav1.8 has been detected in the aging heart (Dybkova et al., 2018) and mutations in Nav1.8 have been associated with cardiac pathologies, although this may be explained by the presence of Nav1.8 in intracardiac neurons (Verkerk et al., 2012). Alternatively-spliced Nav channels add further variations, including an exon-skipping event for Nav1.5 in macrophages (Rahgozar et al., 2013). Further investigation into the sequence- and structure-specific differences in Nav channels may elucidate their functional roles in different tissues.

The revolution in cryogenic electron microscopy (cryo-EM) has revealed unparalleled insights into the structures of the Nav channels and the regulatory β-subunits (Jiang et al., 2022; Blundell and Chaplin, 2021; Liang et al., 2022). High-resolution cryo-EM structures of all human Nav channel α-subunits except Nav1.9 have been published (Noreng et al., 2021). The Nav channel tertiary structure is comprised of four internally homologous domains (DI-DIV) that enclose a central pore in which ion selectivity is determined by size and charge (Figures 1A and 1B) (Catterall, 2000). The pore is surrounded by extracellular turret loops (ECTLs), contributed by each of the domains (Stephens et al., 2015). Helices S1-4 of each domain form the Voltage-Sensing Domain (VSD), in which positively-charged lysine and arginine residues line one face of the S4 α-helix (Catterall, 1986). In response to changes in membrane potential, the movement of the S4 helices in domains I-III induce channel activation, whilst the subsequent movement of the domain IV S4 helix induces inactivation (Angsutararux et al., 2021).

**Figure 1.**
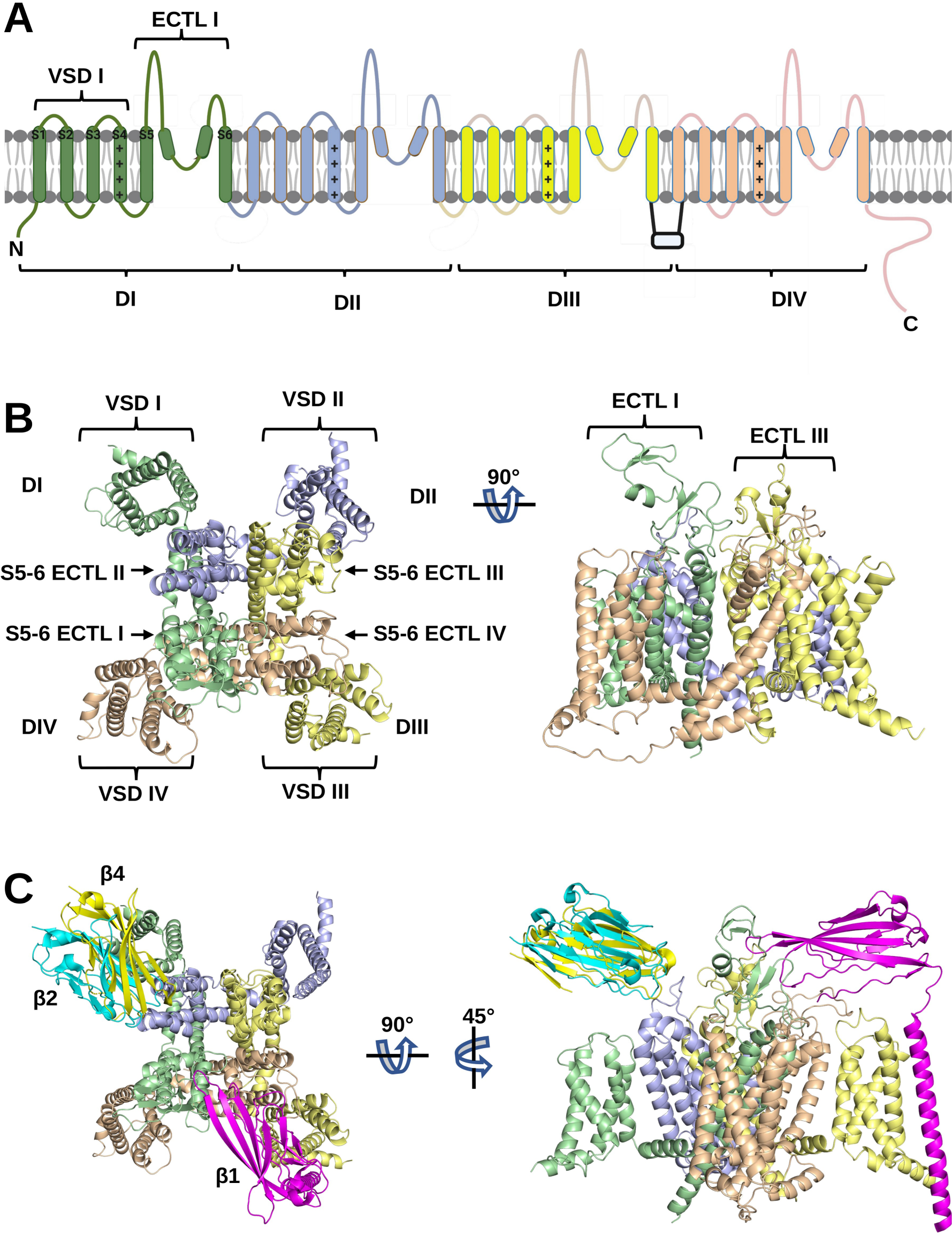
Nav channel α-subunit domains and β-subunits. (A) Key structural features of the Nav channel α-subunit. The four internally homologous domains, DI-IV, are colour-coded and labelled. Within DI, the transmembrane helices, S1–6 (including the positively-charged S4 helix), the voltage-sensing domain (VSD) and the S5-6 extracellular turret loop (ECTL) are labelled. (B) Cryo-EM structure of human Nav1.5 in top and side view (PDB: 7DTC). (C) The resolved binding sites of β1 (magenta), β2 (cyan), and β4 (yellow) based on cryo-EM structures of human Nav1.1 (PDB: 7DTD); Nav1.2 (PDB: 6J8E) and Nav1.4 (PDB: 6AGE), as depicted from top and side views.

Nav β-subunits (β1-4) modulate channel gating, trafficking, and kinetics (Namadurai et al., 2015). The β-subunits comprise a single N-terminal extracellular Ig-domain that is connected via a flexible linker to a transmembrane domain and C-terminal intracellular tail (Salvage et al., 2023b; Cusdin et al., 2010). The interactions between several of the Nav channels and β-subunits have been determined by cryo-EM. Specifically, human β1, β2, and β4 have, respectively, been resolved with Nav1.1 (Pan et al., 2021), Nav1.3 (Li et al., 2022), Nav1.4 (Pan et al., 2018), Nav1.6(Fan et al., 2023), and Nav1.7 (Shen et al., 2019; Huang et al., 2022); Nav1.3 (Li et al., 2022), Nav1.6(Fan et al., 2023), and Nav1.7 (Shen et al., 2019; Huang et al., 2022); and Nav1.1 (Pan et al., 2021). In all of these structures, the β1 transmembrane domain makes extensive contacts with the S1 and S2 helices of the α-subunit domain III VSD, while the β1 Ig-domain contacts extracellular regions in the domain III VSD and the ECTL regions of domains I and IV (Figures 1C and 1D). In contrast, the Ig-domains of β2 and β4 contact the α-subunit domain II ECTL via a covalent disulfide bond, but with no apparent interactions of the β-subunit transmembrane domain. The binding site of β3 is not yet clear. However, the strong sequence similarity between β3 and β1 (Namadurai et al., 2015; Salvage et al., 2023b) suggests that β3 may bind to these Nav channel isoforms in a similar way as β1 (Zhu et al., 2017). Indeed, the recent cryo-EM structure of β3 and the non-classical sodium channel NaX is consistent with this view (Noland et al., 2022). The β-subunits are structurally related to members of the Ig-domain family of cell-adhesion molecules. They have the potential to form trans and cis homo-oligomers that interact with β-subunits on opposing membranes or neighboring plasma membranes, respectively, via their Ig domains: the β1, β2, and β4 have been demonstrated to interact in trans and the β3 subunit has been suggested to interact in cis (Salvage et al., 2020a). Such homophilic interactions may lead to supramolecular clustering between the Nav channels to modulate localized depolarization (Salvage et al., 2020b).

The extracellular domains of plasma membrane proteins often contain one or more copies of the canonical N-linked glycosylation motif: NX[S or T], where X is any amino acid except proline (Gavel and Heijne, 1990). Changes in N-linked glycosylation, e.g. as a result of a mutation, may result in changes in protein folding, intra- and intermolecular interactions, and degradation (Freeze and Schachter, 2009; Beaudoin et al., 2022). In the endoplasmic reticulum, a core glycan structure containing two N-acetylglucosamine (GlcNAc) residues and three mannose residues can be covalently added to the asparagine residue of this motif, provided it is accessible to the glycosylation enzymes (Reily et al., 2019). During further maturation, the outer branches of the tree become heterogeneous, reflecting the stochastic addition and removal of individual sugar residues as the protein traffics through the secretory pathway (Cherepanova et al., 2016). These further modifications may include the addition of one or more sialic acid residues at the terminal positions (Nagae et al., 2020). Nav channel α-subunits contain multiple N-linked glycosylation sites in their ECTLs. Interestingly, the Nav channel cryo-EM data all show the presence of additional electron density around the relevant ECTL asparagine residues, indicating that they are indeed glycosylated. However, usually only the primary N-acetyl glucosamine rings, covalently attached to the asparagine residue, are resolved, since the full glycan trees are normally too flexible to determine single structures (Atanasova et al., 2020). Thus, the existing cryo-EM structures may risk de-emphasising the full extent of the glycan trees and their structural implications. Herein, the structural effects of glycans are questioned for Nav1.5 and other Nav channel isoforms, with particular emphasis on the positioning of sialic acid residues close to VSDs and the potential for steric clashes between β-subunits and the glycosylated α-subunit (Salvage et al., 2023b, 2020a).

## 2. Results

### 2.1. Structural modelling and spatial organization of N-linked glycans on Nav channels

The cryo-EM models of Nav1.1-1.8 are all resolved with at least the first glycan detectable moiety at various NX[S or T] motifs exposed on the ECTLs. These sites provide experimental references for predictions that may uncover additional potential sites. The preliminary use of sequence-based bioinformatics tools (NetNGlyc (Gupta and Brunak, 2002), N-GlycoSite (Zhang et al., 2004)) to predict N-linked glycosylation resulted in several false-negative and false-positives. Therefore, only those sites where an asparagine-proximal N-acetyl glucosamine residue was discovered on the cryo-EM structures were inspected for their structural impact (Figure 2A).

**Figure 2.**
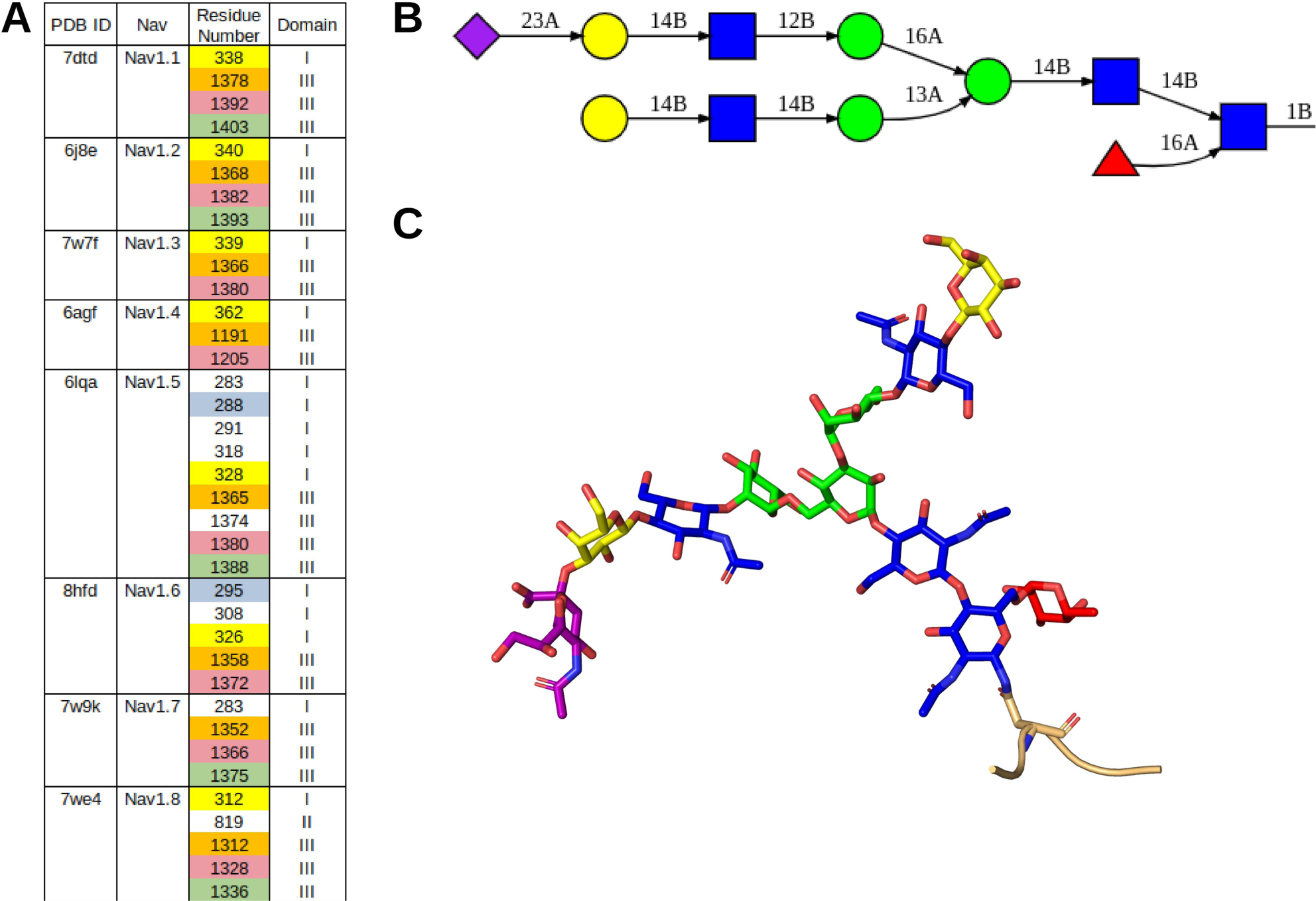
Position and structural representation of Nav glycans. (A) The N-linked glycan positions on Nav channel cryo-EM structures are listed in a table format. Each color denotes homologous N-linked glycosylation sites within the indicated α-subunit isoform. (B) The representative N-linked glycan tree used in this study is shown in 2D format. (C) The corresponding 3D representation of the glycan tree shown in (B). N-acetylglucosamines are shown in blue, mannose in green, galactose in yellow, fucose in red and sialic acid in pink. Connectivity (alpha = A, beta = B) between glycan moieties is notated (B).

Although mature N-linked glycan trees are heterogeneous, they all share a common core sequence of N-acetylglucosamine, mannose and galactose. Most are ‘complex’ with branching antennae. The tips typically contain at least one sialic acid residue. Previous work that is currently under peer review has identified the most prevalent glycan tree on the β3-subunit as a core-fucosylated, bi-antennary structure contain one terminal sialic acid. Hence, this form was used as a representative template for glycan modelling (Figures 2B, C). Alignment of the Nav channel primary sequences with reference to the resolved sites on the cryo-EM structures reveals three sites common to all channels (corresponding to N291, N1365, and 1380 on Nav1.5), one site (N328 on Nav1.5) on domain I shared between all channels except Nav1.7 and Nav1.9, and one site (N1388 on Nav1.5) on domain III shared between Nav1.5, Nav1.7, and Nav1.8 (Figure 3A). Of note, the glycan moieties of some conserved NX[S or T] motifs were not seen in the cryo-EM structures, since these motifs are present in flexible regions of the ECTLs and, thus, were not resolved. Although absent from the resolved structures due to the flexibility of ECTL I, the primary sequence of the Nav1.4 ECTL I shows the highest number of N-linked glycosylation motifs – seven – at this site. Among unique N-linked glycosylation motifs on the α-subunits surfaces, Nav1.5 was found to have four total on domains I and III, Nav1.6 has two on domain I, and both Nav1.7 and Nav1.8 have one on domains I and II, respectively. With the exception of the Nav1.8 N819 residue on domain II, all N-linked glycosylation sites were noted on the ECTLs of domains I and III. The ECTL I is shown to have the highest density and largest variability of N-linked glycan motif start sites.

**Figure 3.**
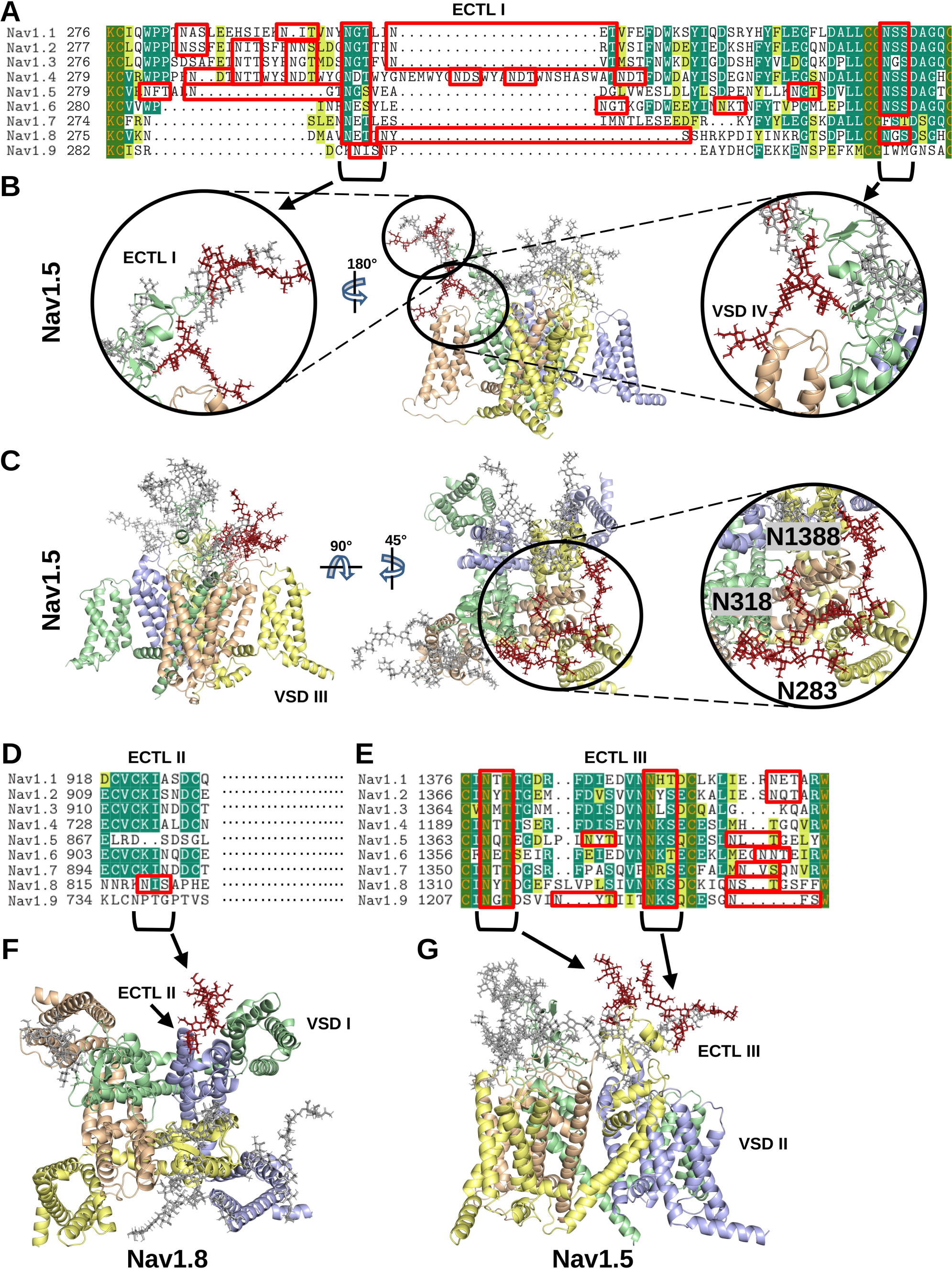
N-linked glycans on Nav channels. (A) Alignments of the amino acid sequences of the Nav1.1-1.9 domain I ECTLs are shown. The residues are colored by similarity (yellow), conservation among 50% of sequences or higher (green), or 100% conservation (orange letters and in green blocks). All NX[S or T] motifs in the sequence alignments are outlined with red rectangles to note evolutionary conservation of the motifs. (B, C) The conserved glycans on domain I ECTLs are colored in crimson and shown on the modelled Nav1.5 structure. In (C), the unique Nav1.5 glycans above VSD III are shown in crimson. (D, E) Alignments of the amino acid sequences of the Nav1.1-1.9 domain II and III respectively. Residue conservation and NX[S or T] motifs are colored as in (A). (F) The unique Nav1.8 N-linked glycan N819 is shown in crimson on the modeled Nav1.8 structure. (G) The conserved glycans on domain III are colored in crimson and shown on the modelled Nav1.5 structure.

Structural modelling of the glycans may provide further insight into the positioning of the glycan trees relative to functional domains on the protein surface. Accordingly, representative glycan trees were modelled at each of the sites resolved on the Nav1.1-1.8 cryo-EM structures. As shown in Figure 3A-G, the common sites found on domain I are located above the VSD of domain IV, and the common sites on domain III are found above the VSD of domain II around the pore. Interestingly, the glycan attached to N328 in Nav1.5 and shared with all Nav channels except Nav1.7 and Nav1.9 presents its negatively-charged sialic acid moiety in proximity to the DIV VSDs (Figure 3C). Two of the three unique glycosylated residues on Nav1.5 (N283, N318, N1388 are positioned above the VSD of domain III (Figure 3C). Interestingly, the increased presence of other glycans on domain I (N283, N288, N328) and domain III (N1365, N1374, N1380) may be sterically inducing a re-direction of the three aforementioned glycans towards the area above the domain III VSD. The unique Nav1.8 N-linked glycosylation site at N819 is positioned near to the VSD of domain I – which is novel among the other Nav channels – and, thus, may interact with the VSD or nearby ECTLs (Figures 3E and 3G). The two conserved glycans on ECTL III are shown to be near the pore (N1365 on Nav1.5) and above VSD II (N1374 on Nav1.5) (Figures 3F and 3H). Further structural dynamics may provide additional insight into the flexibility and, thus, density of the glycans on the Nav channel surface.

### 2.2. Structural effect of Nav channel N-linked glycans on β-subunit accessibility

Nav channels are involved in regulating the action potential of diverse cell types in the nervous and muscular systems. For example, Nav1.4 and Nav1.5 are known for propagating action potentials in skeletal and cardiac muscle, respectively (Loussouarn et al., 2016). Although differences in protein sequence and structure between Nav1.4 and Nav1.5 have been investigated to better understand the physiological differences between muscle types (Salvage et al., 2023a), comparing the differences in glycan conformational landscape may shed light on the functional differences that permit different types of action propagation specific to the tissue. Furthermore, Nav1.4 shares similar glycosylation profiles with Nav1.1, Nav1.2, and Nav1.3 and, thus, may serve as a representative of such structures. On the other hand, Nav1.5 has a much higher density of glycans on its surface and further analysis may provide insights into the conformational landscape of such extensive glycosylation. Therefore, to investigate glycan tree flexibility and interactions, both at sites shared between isoforms and for Nav1.5-specific sites, all-atom molecular dynamics simulations were conducted with the glycosylated and non-glycosylated Nav1.4 and Nav1.5 channels in a representative plasma membrane (Figure 4A-F).

**Figure 4.**
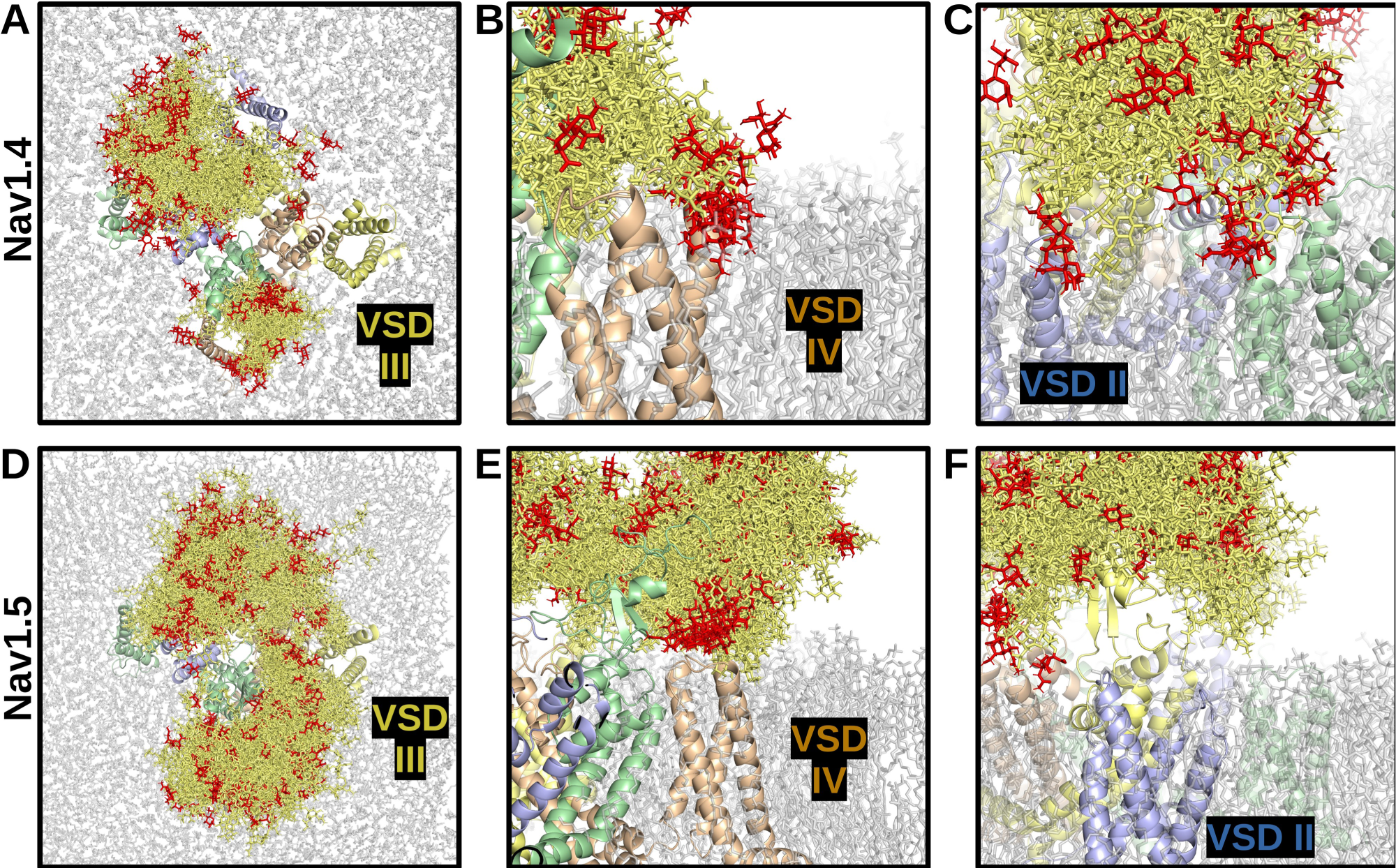
N-linked glycan density on Nav1.4 and Nav1.5. Images show overlapping and superimposed snapshots of glycan conformations extracted from the molecular dynamics simulations every 5 nanoseconds, over a time frame of 150 nanoseconds. (A) Top view of Nav 1.4. (B, C) side views of Nav1.4, VSDs IV and II respectively. (D) top view of Nav1.5. (E, F) side views of Nav1.5, VSDs IV and II respectively. glycans interacting with VSD IV are depicted. The Nav1.4 (C) and Nav1.5 (F) glycans interacting near VSD II are displayed.Glycans are shown in dark gold, with sialylated tips in red.

The molecular dynamics simulations reveal a broad landscape of conformations and interactions displayed by the glycans. Importantly, in both channels, the pore is not blocked by glycans. The lack of density above the channel opening suggests that ion conductance would not be hindered by the presence of glycosylation sites near the pore (e.g. N1191 on Nav1.4 and N1365 on Nav1.5). In contrast, for both channels, the ECTLs of domains I and III are completely covered by glycans on their corresponding loops and the ECTL of domain II is largely overshadowed by the glycans on the ECTL of domain I. The VSDs of domains II and IV were discovered to potentially interact with glycans, which occurred in both Nav1.4 and Nav1.5 α-subunits (Figures 4B and 4E). Notably, in the simulations, the mannose, galactose, and N-acetylglucosamine moieties more than the sialic acid tips remained closer to the basic residues within the VSD throughout the simulations. Based on the simulation data, which shows that the sialic acids interacted with channel residues and membrane lipids surrounding the inner portion of the VSD, the sialic acids may, nevertheless, interact with residues within the VSD and affect gating and kinetics of the channel. The glycans pointing parallel to the membrane from the ECTL of domain III (i.e. N1205 on Nav1.4 and N1380 on Nav1.5) were not found to interact with the VSD of domain II and were more likely found to interact with membrane head groups in the space between the VSDs of domains II and I (Figures 4C and 4F).

The conformations of the glycan trees over the course of the simulations were also compared with known β-subunit binding sites to determine how their positions may affect binding. Here, analysis of the total 3D space occupied by the glycan conformations revealed a notable difference between Nav1.4 and Nav1.5, as depicted in Figures 4A and 4D, respectively. Specifically, the ECTL of domain IV, and the VSM of domain III are exposed in Nav1.4 but are completely covered in Nav1.5. This isoform-specific difference reflects the presence of Nav1.5-unique glycosylation sites at N283 and N318, together with another glycan, N1388, on the domain III ECTL, which primarily points towards the ECTL of domain IV. These domains III and IV regions correspond to the binding site of the β1 Ig-domain in Nav1.4 (Figure 5A, C) (Pan et al., 2018). Currently, there are no cryo-EM structures for Nav1.5 and β-subunits (Li et al., 2021; Jiang et al., 2020). But, mutagenesis evidence suggests that the β1 subunit transmembrane domain binds close to Nav1.5 domain III VSD (Zhu et al., 2017), possibly in a manner similar to its binding in Nav1.3 (PDB: 7w77), Nav1.4 (PDB: 6agf), Nav1.6 (PDB: 8GZ1), and Nav1.7 (PDB: 7w9k). However, unlike these other Nav isoforms, the presence of Nav1.5 glycans at N283, N318, and N1388 will cover the β1 Ig domain binding (Figure 5D, F). Hence, the β1 subunit will still attach to Nav1.5 via its transmembrane domain, but it will not be possible for the Ig domain to bind Nav1.5 in the same way it binds to Nav1.4 (see 2.3, below).

**Figure 5.**
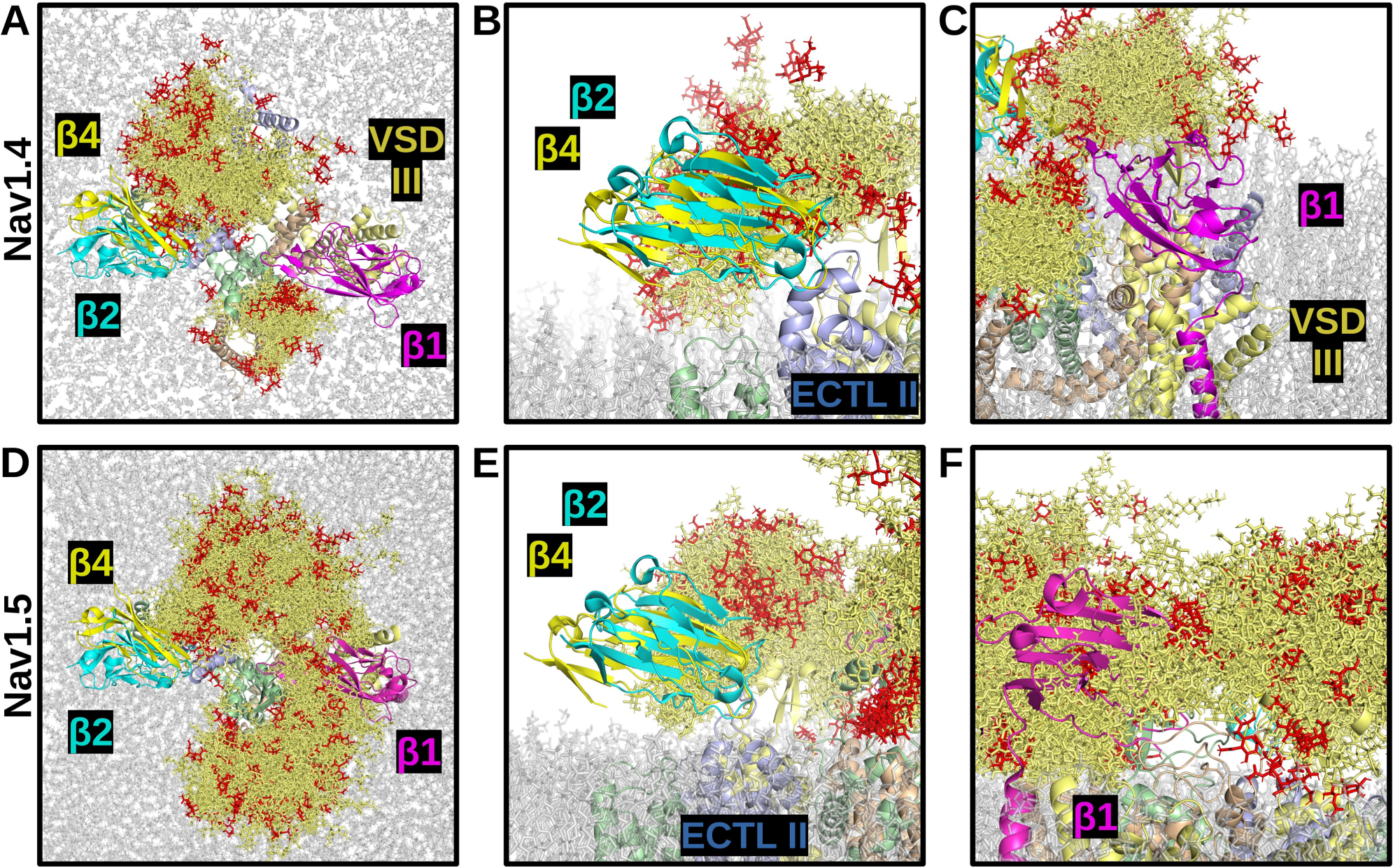
Effect of N-linked glycan density on β-subunit binding for Nav1.4 and Nav1.5. Images show overlapping and superimposed snapshots of glycan conformations from the molecular dynamics simulations, extracted every 5 nanoseconds, over a time frame of 150 nanoseconds. (A) top view of Nav1.4, with β1 (magenta), β2 (cyan), and β4 (yellow). (B) Side view of Nav1.4 interacting with β2, and β4. (C) Side view of Nav1.4 interacting with β1. (D) top view of Nav1.5, with β1, β2 and β4. (E) Side view of Nav1.5 interacting with β2, and β4, assuming the same binding as with Nav1.4. (F) Side view of Nav1.5 interacting with β1, assuming the same binding as with Nav1.4. Glycans are shown in dark gold, with sialylated tips in red.

The VSD of domain I and ECTL of domain II have been shown to be the binding site of β2 and β4 Ig domains (Salvage et al., 2020a). As shown in Figures 5A, 5B, 5D, and 5E, in both Nav1.4 and Nav1.5, the glycans largely do not cover the space at these sites. Thus, glycans will not prevent the binding of the β2 and β4 Ig-domains to Nav1.5. Note, by contrast, that Nav1.8 possesses a unique glycan at N819, which reveals a direct clashing interaction with the putative binding sites of β2 and β4 Ig domains (Figure 3F).

### 2.3. Nav1.5 N-linked glycans may facilitate homophilic interactions between β-subunits

Previous reports have indicated that β1-subunits can interact to permit trans interactions between cells (Salvage et al., 2020b). Our structural analysis of N-linked glycans near the Nav1.5 β1 binding site indicate that the β1 Ig-domain will be unable to interact with the channel. Considering that the β-subunit Ig- and transmembrane domains are connected via a flexible linker, we propose that in Nav 1.5 channels, the β1 Ig domain is less structurally constrained and will be redirected outwards from the channel, which may facilitate the formation of homophilic trans interactions between adjacent membranes expressing Nav1.5/β1.

One case in particular where Nav1.5 and β1 might be engaging in trans homophilic interactions is in the perinexal space of cardiomyocytes (Veeraraghavan et al., 2018). The cardiomyocyte perinexus is a 100-200 nanometer-long anatomically distinct region within the intercalated discs that connects cardiomyocytes and is bounded by connexin-containing gap junctions (Hoagland et al., 2019). Nav1.5 α-subunits associated with the β1-subunit have been shown to cluster on opposing membranes, particularly close to the connexins. It has been suggested that this clustering is facilitated by trans interactions mediated by homophilic binding between mutually opposed β1-subunit Ig domains (Salvage et al., 2023a). The potential structural assembly of such a complex has not yet been explored. Therefore, using protein structure alignment, modelling, and docking methodologies, the structural implications of a Nav1.5/β1 complex interacting in trans with another Nav1.5/β1 complex were explored.

To investigate the structural assemblies that would permit such potential trans interactions, two β1 Ig-domains were, first, docked to one another to find the most likely oligomeric states. Two clusters of dimeric β1 interactions were consistently predicted using different tools: 1) tip-to-tip, wherein the turn regions distal from the transmembrane helix, which are primarily involved in complexing with the Nav channels, are interacting (Figures 6A and 6B), and 2) side-to-side, in which the β-strands interact (Figures 6C and 6D). The predicted binding affinity of the dimer interactions, using PRODIGY (Xue et al., 2016), was −8.1 kcal/mol for the tip-to-tip model and −10.1 kcal/mol for the side-to-side model, suggesting that side-to-side interactions may interact more strongly. The two β1 Ig-domains of both the tip-to-tip and side-to-side models were then pulled apart from the most C-terminal residue, M154, of each flexible linker with steered (constant velocity pulling) molecular dynamics simulations to determine the maximum perinexal intermembane distance based on theoretical biophysical constraints. The β1 dimer structures that were selected to represent the maximum distance were extracted 10 picoseconds before dimer interface rupture. Upon measuring the predicted distance between the lipid head groups, the tip-to-tip model shows a distance of 18.72 nm and the side-to-side model shows 15.34 nm.

**Figure 6.**
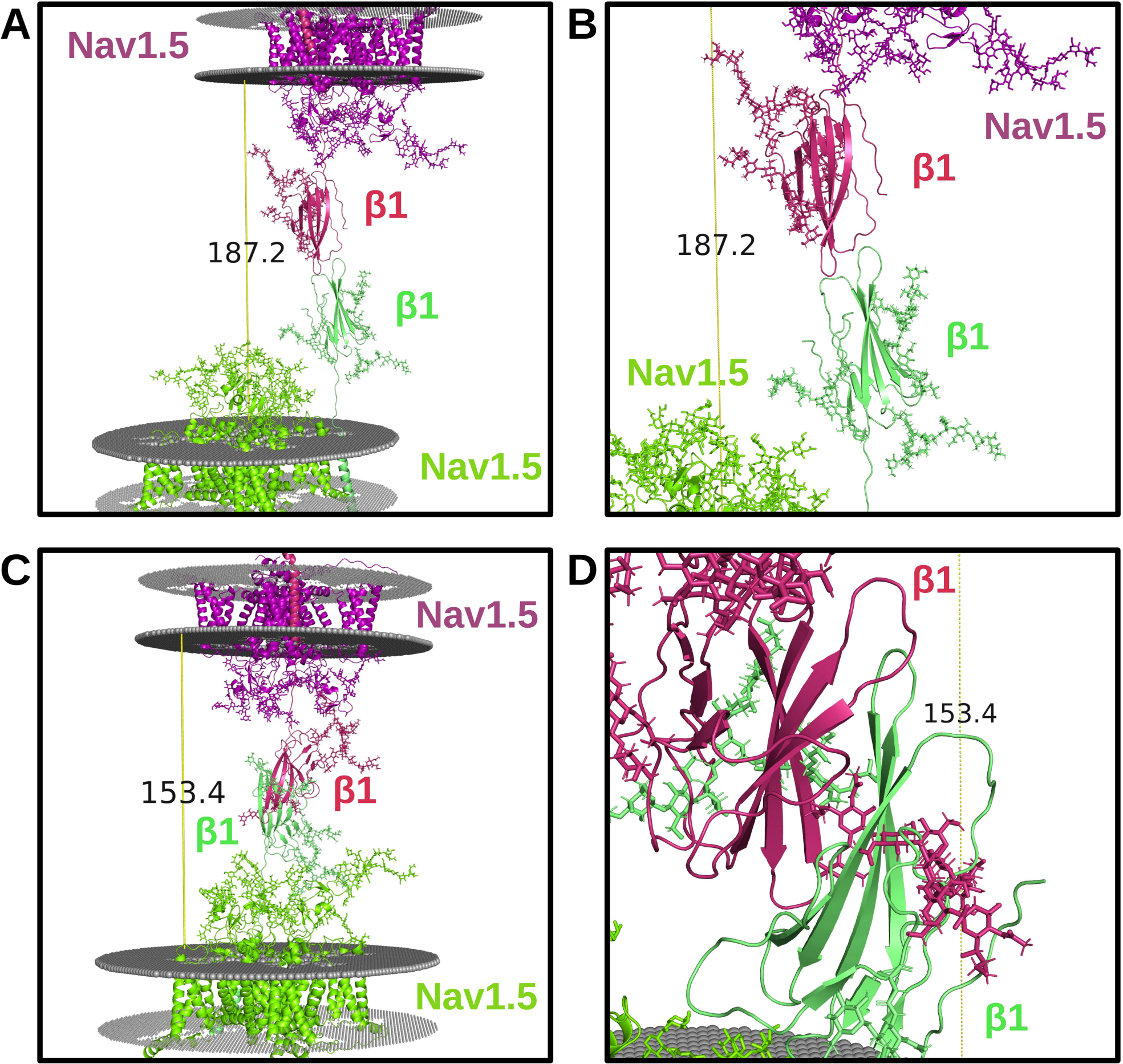
Putative trans interactions between Nav1.5/β1 complexes. (A) Full snapshots and (B) close-up image of the ‘tip-to-tip’ models of the trans interactions between Nav1.5/β1 complexes on opposing membrane (grey). (C) Full snapshots and (D) close-up image of the ‘side-to-side’ models. Note the potential for glycan hindrance in the ‘side-to-side’ model.

Nav β3 has been shown to trimerize and potentially permit homophilic cis interactions (Namadurai et al., 2014). Since the β3 Ig-domain would be redirected in the same manner as β1, both β-subunits may bind at VSD III through interactions between transmembrane domains and create a network of Nav channel supra-clustering on both membranes in the perinexal space. If the transmembrane domains of β2 and β4 do, indeed, interact stably with the Nav1.8 transmembrane domains, cis homophilic dimerization – in which β-strands in the β4 Ig-domains exchange with one another to covalently link the dimer – between β4 subunits may also be relevant in the case of N-linked glycosylation of the ECTL on domain II of Nav1.8. Such dimerization may link Nav1.8 channels together to form higher order structures in the peripheral nervous system. Therefore, N-linked glycans on Nav α-subunits may induce higher-order assembly of Nav channels in specialized membranes that contribute to physiological mechanisms, such as ephaptic conduction (Hoagland et al., 2019).

### 2.4. Clinically-relevant mutations in Nav channel N-linked glycosylation motifs

Further evidence for the functional roles of glycan trees on the Nav channel structures may be revealed by the presence of mutations that both disrupt N-linked glycosylation and manifest into clinical pathologies. Mutations that remove glycosylation sites may occur at the glycosylated asparagine itself, the serine/threonine at the N+2 position, or at the N+1 position if a proline is substituted for the X residue. Thus, disease-causing mutations from the “Disease/Phenotypes and variants section” of each Nav channel UniProt page, an extensive review of Nav variants compiled by Huang et al. (Huang et al., 2017b), and ClinVar (Landrum et al., 2020) were referenced to notate mutations at N-linked glycosylation sites are associated with pathological phenotypes. Only ClinVar entries classified as “Likely Pathogenic” and “Pathogenic” were considered. The mutations are summarized in Figure 7A.

**Figure 7.**
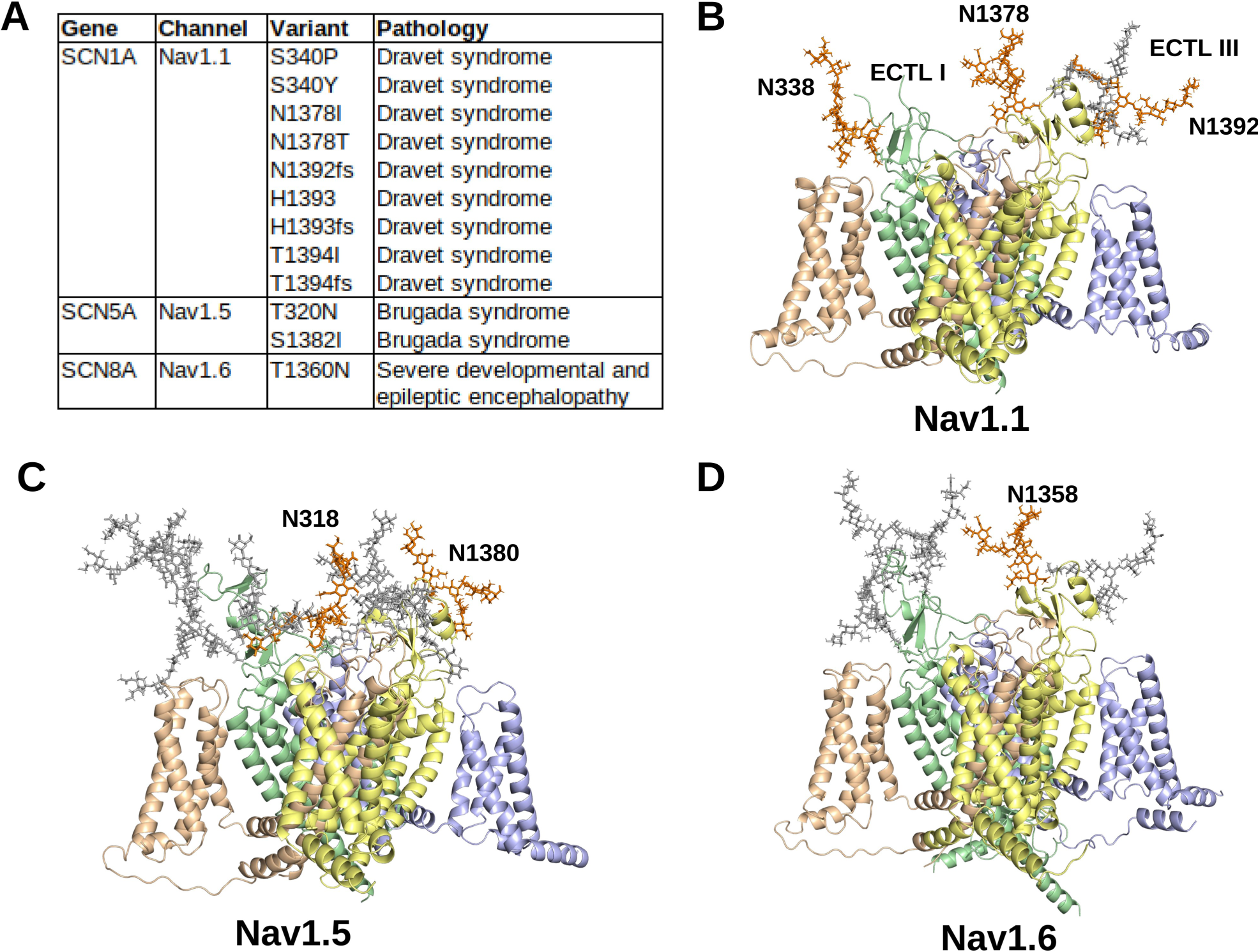
Mutations at N-linked glycan sites linked to clinical pathologies. (A) The N-linked glycan sites disrupted by mutations and their associated pathologies are listed. The N-linked glycans affected by mutations (orange) in (B) Nav1.1; (C) Nav1.5 and (D) Nav1.6 are shown amongst the other N-linked glycans (grey) on the Nav channel surfaces.

Nav1.1 has been discovered to be distributed throughout the central nervous system (Duflocq et al., 2008) and peripheral nervous system (Espino et al., 2022). All four N-linked glycosylation sites – N338, N1378, and N1392 – on Nav1.1 were affected by seven mutations – S340F (Depienne et al., 2009), S340P (Depienne et al., 2009), S340Y (Landrum et al., 2020), N1378H (Zuberi et al., 2011), N1378T (Zuberi et al., 2011), H1393P (Stefanaki et al., 2006), T1394I (Zuberi et al., 2011) – in Dravet Syndrome patients. Three clinically-relevant frameshift mutations – N1392fs, H1393fs, and T1394fs – occurred at the N1378 site and were associated with Dravet Syndrome, but the recorded clinical outcomes likely stem from downstream effects of the frameshift mutations on protein structure that are unrelated to glycosylation. The mutations at S340 and H1393/T1394 may prevent interactions between the glycan and the VSDs of domains IV and II, respectively, which brings further attention to the putative interactions between the negatively-charged glycan sialic acids and the positively-charged VSD residues. The N1378H and N1378T variants may disrupt incorporation of the glycan located above the pore, which may affect several processes, such as gating, trafficking, or protease inhibition. These mutations (Figure 7B) affect all but one glycosylation site on Nav1.1, which reveals the selective functional roles for glycosylation in regulating neural Nav channel activity.

Nav1.5 is associated with cardiomyocyte action potential propagation (Rook et al., 2012). Two sites – N318 and N1380 – were affected by two mutations – T320N (Kapplinger et al., 2010) and S1382I (Kapplinger et al., 2010; Smits et al., 2002) – on Nav1.5 in Brugada Syndrome patients (Figure 7C). The T320N mutation removes the glycan that is centrally positioned among the other glycans for preventing interactions with the β1 and potentially β3 Ig-domains. These data potentially provide further evidence that blocking the β-subunit Ig-domains from binding to the channel is vital for cardiomyocyte function. The Nav1.5 S1382I mutation may prevent interactions from the glycan that is located above the VSD of domain II. These data further add to the notion that glycans may interact and modulate the structure-function relationship of VSDs.

Nav1.6 is widely expressed throughout the brain and regulates the initiation of action potentials at nodes of Ranvier (Wagnon et al., 2016, 10). Only one mutation – T1360N – was found to interfere with N-linked glycosylation and was associated with global developmental delay and seizures (Figure 7D) (Wengert et al., 2019). Similar to the mutations at N1378 in Nav1.1, the Nav1.6 T1360N variant may prevent glycosylation above the pore, which could affect ion conduction and gating.

The presence of variants associated with human pathologies at N-linked glycosylation sites provides additional support for the critical roles of such glycans in the structure and function of Nav channels. Mutations at specific N-linked glycans may also shed light on the importance of the position of the glycans on the Nav channel surfaces. Among the three sites conserved by all Nav channels, mutations in all three were independently associated with a clinical pathology in Nav1.1, and only one conserved site was affected by a mutation in pathologies related to Nav1.5 (T320) and Nav1.6 (T1360). The other recorded clinically-relevant Nav1.5 mutation is unique to Nav1.5 and, as reported above, may affect interactions with the β1 and β3 subunit Ig-domains. Of note, synonymous mutations at N-linked glycosylation sites were reported in Nav1.3, Nav1.5, Nav1.6, Nav1.7, and Nav1.8 in the ClinVar data, all of which were classified as “Benign” or “Likely Benign”. Further investigation into the clinical relevance of Nav channel N-linked glycans may provide mechanistic insights into their functional roles, and biophysical experimentation of the effect of N-linked glycans on Nav channels may guide clinical and give rise to novel diagnostic and therapeutic options.

## 3. Discussion

Glycosylation of Nav channels has been implicated in altering biophysical properties, such as gating, trafficking, and localization (Zhang et al., 1999; Cortada et al., 2019; Ednie and Bennett, 2012; Wang et al., 2021). Although several studies have outlined the effects of differing composition and amounts of glycosylation *in vitro* and *in vivo*, little work has been conducted regarding the structural effect of N-linked glycans on Nav channels and their impact on interactions between the channels and accessory β-subunits. Therefore, herein, the potential interactions, conformational landscape, and clinical effects of N-linked glycans on Nav channels extracellular surfaces were investigated using sequence and structure-based bioinformatics methodologies.

Sequence and structure comparisons revealed three glycan sites that were found to be common to all Nav channels, which are located on the ECTL of domains I and III. Interestingly, two of the three sites are positioned directly above the VSDs of domains II and IV. Glycan sialic acid residues are known to influence the electric field sensed by the VSDs (Ednie and Bennett, 2012). Since negatively-charged sialic acid moieties have been documented to be present on the termini of glycan tree branches, the voltage-sensing properties of the channels may be altered due to interactions between the positive residues in the VSD and glycans (Robinson et al., 2023). The remaining glycan is found nearly above the channel pore on the ECTL of domain III. However, since the glycan is directed outwards from the channel, it may function to modulate or prevent interactions with other proteins. Interestingly, in examining the sites on cryo-EM structure of the electric eel Nav1.4, which is the only other available resolved non-mammalian vertebrate Nav channel structure, all three conserved human N-linked glycosylation sites are the only sites present (Yan et al., 2017). These data point to deeply evolutionarily conserved functional roles for glycosylation at these three sites.

Nav1.5 was recorded to contain four unique motifs, which were widely distributed on the ECTLs of domains I and III. Notably, two glycans (N283 and N318) on domain I and one glycan (N1388) on domain III were found to largely occupy the area above the VSD of domain III, which, thus, directly covers the binding site of the β1 Ig-domain. The unique glycans on domains I and III of Nav1.5 may, thus, prevent the β1 Ig-domain from interacting with the channel. Although the Ig domain is itself rigid, it is connected to the transmembrane helix via a flexible, disordered neck (Salvage et al., 2020a). Hence, a β1-subunit bound to a fully glycosylated Nav1.5 α-subunit via its transmembrane helix, will impart a greater degree of conformational flexibility on the Ig domain than would otherwise be the case. One example where this may be physiologically important is within the opposing perinexal membranes of the intercalated discs between adjacent cardiomyocytes, where Nav1.5 and β1-subuntis are tightly packed (Veeraraghavan et al., 2018). The perinexal space between these membranes has been shown to facilitate ephaptic conduction between cardiomyocytes (Hichri et al., 2018). Mathematical modelling studies have revealed that the distance between membranes is critically important for ephaptic conduction, no more than about 20nm is optimal (Lin and Keener, 2010; Mori et al., 2008). The β1-subunit Ig domain is known to be capable of trans-mediated cell-adhesion, and inhibition of β1-mediated trans adhesion within the perinexus leads to reduced ephaptic conduction (Veeraraghavan et al., 2018). The distance between the membranes that define the perinexal space is about 15-20 nanometers (Veeraraghavan et al., 2018). But, in many places, particularly where closer to the connexin gap junctions, this distance is less (5-10 nm). Our modelling identified two possible modes of trans homophilic binding between β1-Ig domains: ‘tip to tip’ or ‘side by side’ with estimated membrane-to membrane distances of 18.72 nm and 15.34 nm respectively (Figure 6). But, these are maximum estimates, assuming the flexible neck connecting the Ig- and transmembrane domains is fully extended. Thus, both models are consistent with the morphological data. An inhibitory peptide was described by Veeraraghavan *et al*., which successfully disrupted the trans homophilic binding between β1 Ig domains, would be expected to interfere with a ‘side to side’-type interaction between Ig domains (Veeraraghavan et al., 2018). However, the β1 Ig-domain contains a glycan at N135 and in the side-to-side model, this is pointing directly into the other β1 Ig-domain (Figure 6). This would make a ‘side to side’ model of the type identified here highly unlikely. Thus, the ‘tip-to-tip’ model may be more likely to represent a stable state for trans interactions. As further support, a ‘tip-to-tip’ model has also been proposed for cis homophilic interactions between β3 subunits (Glass et al., 2020).

The analysis also identified a N-linked glycan unique to Nav1.8 which would occlude the binding sites of the β2 and β4 Ig-domains. Therefore, such glycosylation may either prevent interactions with the β-subunits entirely – since the transmembrane interactions are likely to be more transient than β1 as noted by their absence in resolved cryo-EM structures – or the Ig-domains may be redirected as in the proposed case of β1 and potentially β3. It should further be noted that Nav1.8 and Nav1.5 both lack a free cysteine residue on domain II ECTL and thus cannot form the covalent disulfide bond with the β2 and β4 Ig domains that occurs with most Nav channels (Salvage et al., 2020a). Interestingly, the β4 Ig domain can form cis-interacting covalent dimers using the cysteine residue that forms a disulfide bond to most Nav α-subunits (Shimizu et al., 2017). Hence, the absence of a suitable cysteine residue on Nav1.5 and Nav1.8, together with a blocking glycan on Nav1.8, would further facilitate cis-interactions between β2 and β4 (Salvage et al., 2020a).

The β-subunits affect the electrophysiological properties of the Nav channel α-subunits (Namadurai et al., 2015). Hence, these data may have important implications for understanding Nav 1.5 and 1.8 channel gating (Vijayaragavan et al., 2004; O’Malley and Isom, 2015). Distinct glycosylation states of Nav channel α-subunit isoforms, including Nav1.5, have been detected (Mercier et al., 2015; Laedermann et al., 2013). N-linked glycosylation and Nav channel oligomerization occur contemporaneously within the endoplasmic reticulum at either independent or co-dependent rates (Braakman and Hebert, 2013). Glycosylation of Nav1.5 α-subunit at N318 and N283 will prevent the binding of the β1 Ig domain to DIII VSD, but not its association with the α-subunit via its transmembrane helix. But if the Nav1.5 α-subunit assembles with β1 before it is glycosylated, would that permit the binding of the β1 Ig domain to the DIII VSD? If so, then a cell co-expressing Nav1.5 α- and β1-subunits may contain a mixed population of Nav1.5/β1 hetero-oligomers with distinct structural properties and perhaps distinct functional behaviour (Schoberer et al., 2018; Ninagawa et al., 2021). Their proportion would depend on the relative rates of glycosylation vs hetero-oligomer assembly. The Nav channel glycosylation patterns and rates may also be dependent on the expression of specific glycosyltransferases and, thus, may be cell or tissue specific (Medzihradszky et al., 2015). Further work is required to underpin the competition dynamics between β-subunit binding and glycosylation of Nav channel α-subunits within the endoplasmic reticulum.

The conformational variability of the Nav channel glycans creates a notable shield around the Nav channel surfaces (Seitz et al., 2020). In addition to altering the binding sites of β-subunits, glycans on Nav channel surfaces may interfere with binding to toxins, proteases, and other proteins (Beaudoin et al., 2022). Additionally, consideration of the glycan structural conformational space may guide studies targetting Nav channels with inhibitory or activating small molecules, peptides, and antibodies (Seitz et al., 2020). For example, the spider toxin Dc1a binds to the DII VSD of Nav1.7, which is in agreement with the data described above considering that the DII VSD is largely unoccupied by glycans (Bende et al., 2014). Furthermore, incorporating isoform-specific glycosylation structural information may further inform the targeting of specific Nav channels. For example, the glycan overshadowing the DIV VSD of Nav1.5 (N328) is conserved among Nav channels with the exception of Nav1.7 and Nav1.9. Indeed, the DIV VSD of Nav1.7 has been shown to confer isoform-specific targeting of Nav1.7 and is also the binding site of venom toxins (e.g. OD1) specific to Nav1.7 (Kschonsak et al., 2023; Salvage et al., 2023c; Jalali et al., 2005).

Therefore, alongside the electrostatic and structural effects, further investigation into the roles that glycans play in modulating interactions between Nav channels and other binding partners may reveal new insights into higher order complexes at the cell surface, coordination of intracellular signaling pathways, and site-specific drug target selection.

## 4. Conclusions

Herein, molecular modelling and all-atom molecular dynamics simulations were applied using the resolved Nav channel α-subunit structures and the resolved sugar moieties as references. In particular, a comparative analysis of the skeletal muscle-specific channel Nav1.4 and the heart muscle-specific channel Nav1.5 cryo-EM structure sites was conducted to better understand the landscape of glycan conformations with respect to Nav channel domains and β-subunit binding. Molecular dynamics simulations revealed that negatively-charged sialic acid residues of two conserved glycans may interact with the VSDs of domain IV and II. Notably, three of the five Nav1.5 isoform-specific N-linked glycosylation sites cover the landscape above domain III where the β1 (and likely β3) Ig-domains bind in other Nav channel isoforms. These glycans will thus prevent the binding of the β1 and β3 Ig-domains, allowing them to be redirected outwards and to rotate more freely, while preserving transmembrane domain interactions with the domain III VSD.

It was also noted that Nav1.8 contains a unique N-linked glycosylation site on domain II ECTL that likely prevents interactions with the Ig-domains of β2 and β4. Previously determined complexing between the β1 transmembrane domain and Nav channel VSD III transmembrane domains suggests that β1 and β3 may, nevertheless, bind to Nav1.5 although the Ig-domain would be directed outwards from the channel. Protein-protein docking revealed that this blocked interaction may, thus, redirect the Ig-domain outwards for more likely interactions with other β3 Ig-domains to permit cis supra-clustering of channels or the β1 Ig-domains for trans interactions between channels on opposing plasma membranes, such as those in the cardiomyocyte perinexal space. Further experimental work is necessary to validate these hypotheses. We propose that the isoform-specific structural features of Nav1.5 and Nav1.8 may have evolved to facilitate functional interactions, that would include the promotion of β-subunit-induced trans and cis cross-linking.

## 5. Methods

### 5.1. Sequence accessions and analysis

The amino acid sequences of Nav1.1 (UniProt Accession: P35498), Nav1.2 (Q99250), Nav1.3 (Q9NY46), Nav1.4 (P35499), Nav1.5 (Q14524), Nav1.6 (Q9UQD0), Nav1.7 (Q15858), Nav1.8 (Q9Y5Y9), Nav1.9 (Q9UI33) were retrieved from UniProt (The UniProt Consortium, 2023). Multiple sequence alignments were constructed using the ClustalW algorithm (Thompson et al., 1994) implemented in the R (version 4.1.2) package “msa” (Bodenhofer et al., 2015).

### 5.2. Protein structure, glycan, and membrane modelling

The experimentally resolved structures of Nav1.1-Nav1.8 were downloaded from RSCB PDB (Burley et al., 2023): Nav1.1 (PDB ID: 7DTD), Nav1.2 (6J8E), Nav1.3 (7W7F), Nav1.4 (6AGF), Nav1.5 (6QLA), Nav1.6 (8FHD), Nav1.7 (7W9K), Nav1.8 (7WFW). The NX[S or T] motifs resolved with at least one glycan moiety were considered for glycan modelling. Protein structures were visualized using PyMol (Schrödinger, LLC and DeLano, 2020).

A representative N-linked glycan tree structure (supporting data under peer review) was modelled onto each NX[S or T] motif using the CHARMM-GUI Glycan Reader and Modeller (Park et al., 2019). The glycosylated models of Nav1.4 and Nav1.5 were placed in a representative mammalian cell membrane, as outlined by Ingólfsson et al. (Ingólfsson et al., 2017, 2014), using PPM 2.0 (Lomize et al., 2012) and the CHARMM-GUI Membrane Builder (Lee et al., 2019). The lipid bilayer membrane is comprised of DSM (Upper leaf: 21.0%, Inner leaf: 10.0%), POPC (35.0%, 15.0%), DOPC (3.5%, 1.5%), POPE (5.0%, 20.0%), DOPE (2.0%, 5.0%), POPS (0%, 15.0%), POPI (0%, 5.0%), POPA (2.2%, 0%), CHOL (31.3%, 28.5%). The individual modelled lipids may be visualized in the CHARMM-GUI Archive - Individual Lipid Molecule Library (https://www.charmm-gui.org/?doc=archive&lib=lipid) (Jo et al., 2009).

To model the Nav1.5/β1 trans interactions, the Nav1.5 structure (PDB ID: 6QLA) inserted into the membrane described above was aligned to the Nav1.4/β1 structure (PDB ID: 6AGF) and the linker region connecting Ig-domain to the transmembrane domain of β1 was reoriented towards an opposing membrane using Foldit Standalone (Kleffner et al., 2017). Protein-protein docking of the β1 Ig-domains was performed with ZDOCK (Pierce et al., 2014) and HADDOCK (Dominguez et al., 2003; van Zundert et al., 2016) using default settings to predict dimerization states.

### 5.3. Molecular dynamics simulations

Conventional and steered all-atom molecular dynamics simulations were prepared and run using GROMACS 2021.3 (Abraham et al., 2015) implemented with the University of Cambridge High Performance Computing resources. Periodic boundary conditions were established in a 150×150×150 Å cubic box. Each system was solvated in 150 mM NaCl with a zero net charge using the TIP3P model (Joung and Cheatham, 2008) and were run at 310 K (Bondi, 1964). The Particle-mesh Ewald method was used to calculate long-range electrostatic interactions, and the cutoff for Coulomb interactions and van der Waals interactions were set to 10 Å (Darden et al., 1993). The LINCS algorithm was used to constrain molecular bonds (Hess et al., 1997). All systems were subjected to steepest descent minimization, upon which six series of a 125 picosecond NPT equilibration ensemble with temperature coupling using velocity rescaling (Bussi et al., 2007) and pressure coupling using the Parrinello-Rahman method were conducted (Parrinello and Rahman, 1981). All simulations were run using the CHARMM36 all-atom force field (C36 FF) with 2 femtosecond time steps (Huang et al., 2017a). Duplicate conventional productions of 150 nanoseconds each were run for the glycosylated Nav1.4 and Nav1.5. Steered molecular dynamics simulations were performed using constant velocity stretching by applying an external force to the M154 residue of both β1 subunits (Lu and Schulten, 1999). A virtual harmonic spring attached to the selected residues were pulled at a constant velocity of 5 nm/ns with a force constant of 100 kJ/mol nm^2^ in opposite directions (Guzmán et al., 2008). One steered production replicate was performed for the side-to-side and tip-to-tip models each (Sheridan et al., 2019). VMD was used to visualize snapshot structures from the productions (Humphrey et al., 1996). Snapshot structures were extracted every 5 nanoseconds from the resultant trajectory files and overlapped onto the respective starting glycosylated membrane-bound Nav1.4 and Nav1.5 structures to represent glycan flexibility and conformational landscape over the course of the simulations (Figures 4 and 5) (Pronk et al., 2013).

## Acknowledgements

We greatly thank the team at the University of Cambridge High Performance Computing Centre. SCS was supported by the British Heart Foundation (PG/19/59/34582 to SCS, CLHH, and APJ). SJA was funded by the Biochemical Society Summer Vacation Studentship.

